# Deciphering the role of germline complex *de novo* structural variations in rare disorders

**DOI:** 10.1101/2024.04.03.587925

**Authors:** Hyunchul Jung, Tsun-Po Yang, Susan Walker, Petr Danecek, Isaac Garcia Salinas, Matthew D C Neville, Helen Firth, Aylwyn Scally, Matthew Hurles, Peter Campbell, Raheleh Rahbari

## Abstract

*De novo* structural variants (dnSVs) have emerged as crucial genetic factors in the context of rare disorders. However, these variations often go undiagnosed in routine genetic screening practices. To shed light on their significance in rare disease, we conducted a comprehensive analysis of the largest cohort of parent-offspring whole-genome sequencing data from the UK 100,000 Genomes Project. Our study encompassed a vast cohort of 12,568 families, including 13,702 offspring affected by rare genetic diseases. We identified a total of 1,872 dnSVs, revealing that approximately 12% of the probands harboured at least one dnSV, of which 9% were identified as likely pathogenic in affected probands (151/1696). Advanced parents’ age was found to be associated with an increased chance of having dnSVs in probands. We discovered 148 clustered breakpoints resulting from a single event. 60% of these complex dnSVs were classified into 9 major SV types, and could be observed in multiple individuals, while the remaining 40% were private events and had not been previously reported. We found 12% of pathogenic dnSVs are complex SVs, emphasising the critical importance of thoroughly examining and considering complex dnSVs in the context of rare disorders. Furthermore, we discovered an enrichment of maternal dnSVs at subtelometric, early-replicating regions of chromosome 16, suggesting possible sex-specific mechanisms in generation of dnSVs. This study sheds light on the extent of diversity of dnSVs in the germline and their contribution to rare genetic disorders.

## Introduction

Structural variants (SVs), defined as genetic changes ≥50bp that encompass copy number variants (CNVs)^1^, rearrangements, and mobile element insertions, play an important role in cancer when occurring in somatic cells^2^. They also arise in the germline, with *de novo* structural variants (dnSVs) contributing to rare disorders^3–10^. For instance, chromosomal microarray (CMA), which is capable of detecting submicroscopic CNVs, demonstrated an average diagnostic yield of 12.2% in patients with developmental and intellectual disorders^11^. Beyond CNVs, other types of SVs such as complex SVs involving clustered breakpoints originating from a single event^3^, have provided insights into rare disorders^12,13^, surpassing the explanatory power of CNVs alone. Nevertheless, in contrast to *de novo* single-nucleotide variants, the prevalence and characteristics of dnSVs, particularly complex ones, remain less elucidated in rare disorders, largely due to the substantial technical challenges^14^.

One prominent difficulty arises from the inherent limitations of short-read technologies in accurately capturing and characterising large-scale genomic rearrangements. The restricted read lengths often result in fragmented or incomplete representations of complex structural variations, leading to difficulties in assembling the complete picture of genomic architecture. This issue is particularly pronounced in regions with high sequence similarity, where distinguishing between homologous sequences proves significant computational and analytical hurdles.

Long-read sequencing mitigates the challenges associated with short-read platforms, by offering a more direct span across SVs, thereby enabling better resolution and a more complete representation of complex genomic variations. While long-read sequencing offers unique advantages in studying SVs, the lack of a substantial long-read sequence from the rare disorder cohort highlights the ongoing importance of precise short-read-based SV analytical pipelines^15–17^. These pipelines, essential for detecting a broad spectrum of SVs, play an important role in minimising false positives. This is particularly pertinent in the absence of large cohort population dataset, which hampers accurate filtering and necessitates robust short-read analytical approaches^18^. Consequently, leveraging large-scale short-read sequence datasets with rigorous analytical approaches remains key for a nuanced understanding of SVs in diseases, particularly rare disorders.

To shed light on their significance in rare diseases, we analysed dnSVs identified in 13,702 whole-genome-sequenced parent–child trios from 12,568 families from the rare disease programme of the 100,000 Genomes Project^19^ using rigorous analytical approaches. The rare disease cohort includes individuals with a wide range of diseases, including neurology and neurodevelopmental (NN) disorders that account for half of the cohort, ultra-rare disorders, ophthalmological disorders, renal and urinary tract disorders, cardiovascular disorders, endocrine disorders, and others (**Extended Data Fig. 1a**).

## Results

### Rate of *de novo* SVs and parental age and sex bias

We developed a rigorous pipeline to analyse an average of 13,980 candidate variants (standard deviation = 2,550) per proband, already called by Genomic England using Manta caller ^20^. We identified a total of 1,872 high-confidence dnSVs, all of which were visually inspected (**Methods**). Some of these dnSVs could be validated using previously identified dnSVs called from independent sequencing data on an overlapping set of families. The validation rate was 100% (n=44): 37 candidate dnSVs were confirmed by array/exome-seq from the Deciphering Developmental Disorders (DDD)^21^ study and 7 candidate dnSVs were confirmed using long-read sequencing data from Genomic England (GEL)^19^, respectively (**Methods, Extended Data** Fig. 2).

Using 1,872 high-confidence dnSVs we estimated an overall mutation rate of 0.13 events per genome per generation, in line with previous reports^22–24^ (**Extended Data Fig. 1b)**. The rate of dnSVs varies across the rare disorder categories; such that probands with neurology and neurodevelopmental (NN) disorders and those with cardiovascular disorders exhibit the highest dnSV rate (0.15 event per genome), whereas probands with ophthalmological and hearing and ear disorders show the lowest (0.1 event per genome; **Extended Data Fig. 1c)**. It is worth noting that the rate of dnSVs is marginally higher in the probands (0.13 event per genome) than in unaffected siblings (n=207; 0.09 event per genome; *P* = 0.05). Approximately 12% (n=1,696) of the probands harboured at least one dnSVs. We identified 17 individuals (z-score > 3) with a considerably higher number of dnSVs (n ≥ 3). These individuals, recruited under different rare disease categories, are not among the previously reported germline SNV hypermutators^25^ in this cohort and have no known history of parental exposure to chemotherapy. We found a statistically significant positive correlation between the number of dnSVs and *de novo* SNVs/indels (*P* = 3.87E-07; **Extended Data Fig. 3a**), which is partly explained by the parental age effect^24,26^. However, the mechanistic basis of this correlation remains yet unclear.

Interestingly, we observed a greater enrichment of dnSVs in probands without diagnostic SNVs/indels compared to those with diagnostic SNVs/indels (*P* < 5.00E-02; **Fig. 1a**), suggesting that a significant proportion of unsolved cases is likely to be explained by dnSVs. We also found parental-age effects on the occurrence of dnSVs (**Fig. 1b**, *P*__*paternal*_ < 5.00E-02 and *P__maternal_* < 1.00E-03). Overall, we identified a significant increase in parents’ age at birth in probands with dnSVs compared to those without them (*P* < 5.00E-02). Among the rare disorder classes, a significant difference in parental age distribution is observed in dysmorphic and congenital abnormality syndromes and skeletal disorders (*P* < 0.05), while skeletal disorders remain significant after multiple testing correction (Benjamini-Hochberg corrected *P* < 0.05; **Extended Data Fig. 3b**). Additionally, we observed 67.8% of the phased dnSVs originating from paternal germ cells (**Fig. 1c**), in agreement with previous findings that the majority of germline *de novo* mutations are paternal in origin^27^.

**Figure 1.**
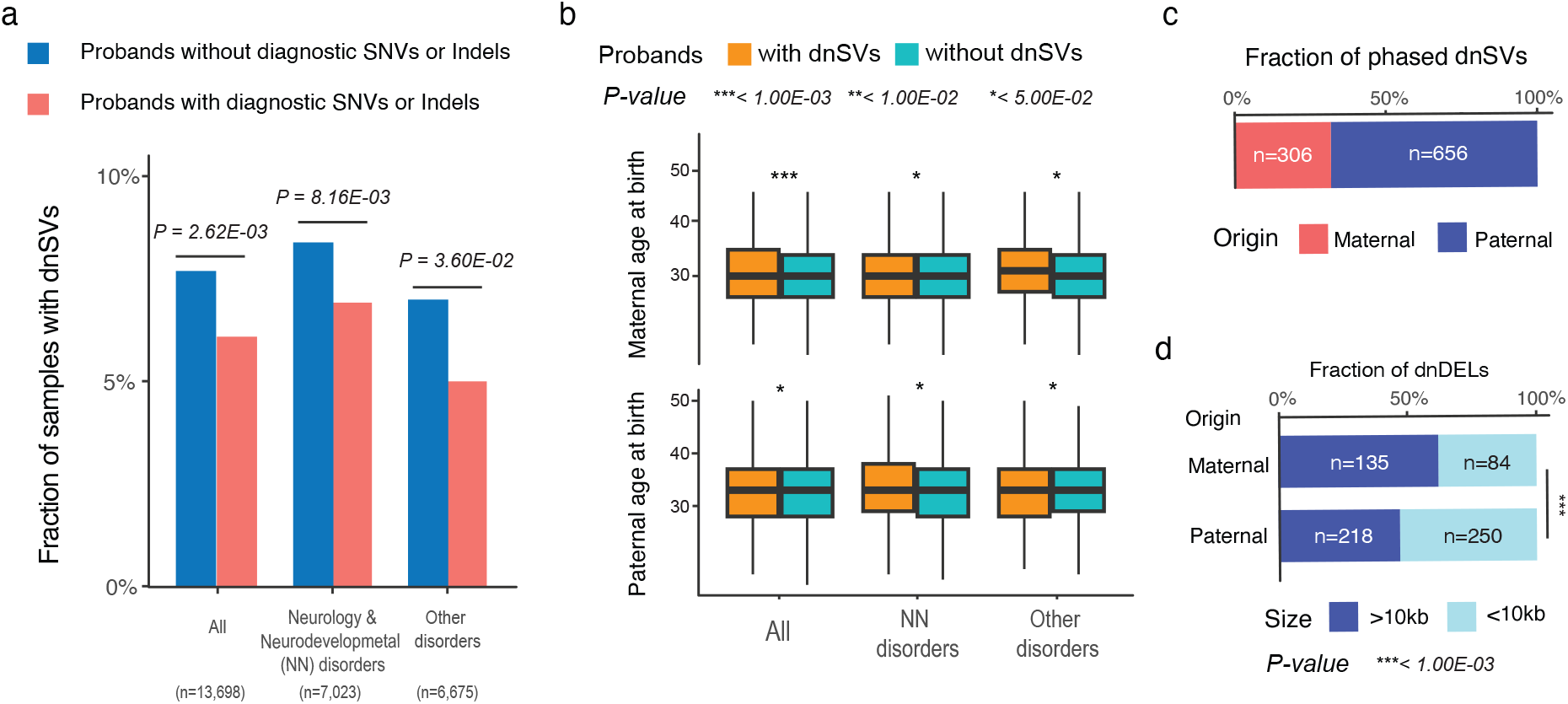
Summary of the identified dnSVs in the rare disease programme of the 100,000 Genomes Project**. (a)** Comparison of the proportion of probands with dnSVs between those with diagnostic SNVs and/or indels and without them. The P values were calculated based on a two-sided Fisher exact test. (**b**) Comparison of maternal and paternal age at birth between probands with dnSVs and those without dnSVs. The P values were calculated based on a one-sided t-test. (**c**) Fraction of phased dnSVs. (**d**) Comparison of the size of dnDELs according to parent of origin. The P value was calculated based on Fisher’s exact test.

### Distribution of different classes of dnSVs

Among the different classes of dnSVs (**Fig. 2a**), simple deletion (n=1,377; 73.6%) was the most common, followed by duplication (n=255; 13.6%), The majority of these duplications are tandem duplications (n=245; 96.5%), followed by dispersed (n=8; 2.7%) and inverted duplications (n=2; 0.8%; **Fig. 2b**). The median detected deletion and tandem duplication sizes are 3.7 kb (range 52 bp - 61 mb) and 49 kb (range 135 bp - 154 mb), respectively (**Extended Data Fig. 3c**). Additionally, we found that maternal dnSVs were enriched for larger deletions (>10 kb), while paternal dnSVs were enriched for smaller deletions (<10 kb) (*P* = 3.04E-04; **Fig. 1d**). We observed a similar enrichment pattern in an independent dataset^24^ (*P* = 6.50E-03; **Extended Data Fig. 3d**).

**Figure 2.**
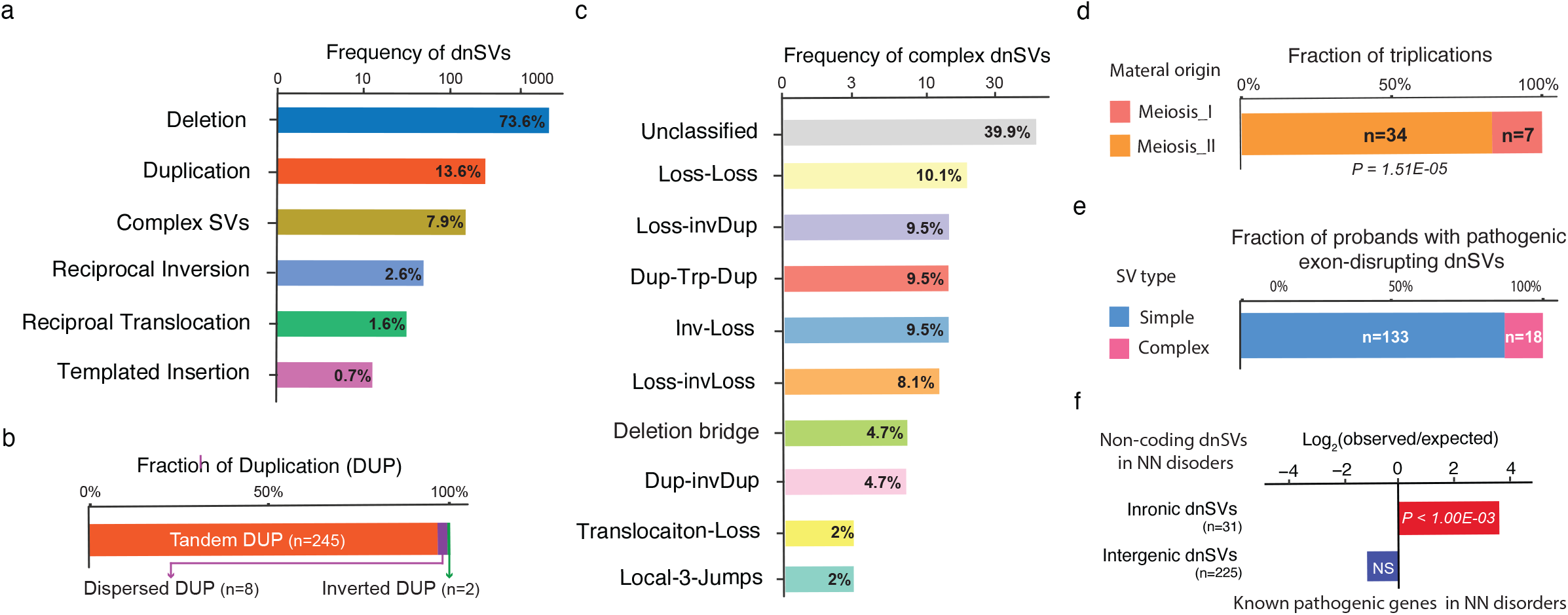
Frequency of dnSVs classes and clinical relevance of dnSVs. (**a**) Frequency of dnSVs classes. (**b**) Subclasses of duplication. (**c**) Frequency of complex dnSVs classes. (**d**) Timing of triplications from maternal origin. Fraction of triplications according to the timing. The p-value was calculated using the exact binomial test. (**e**) Fraction of probands with pathogenic gene-disrupting dnSVs per dnSV type. (**f**) Over-representation of non-coding dnSVs in known pathogenic genes in NN disorders. A window of 50kb up- and down-stream was added to each intergenic dnSV for this enrichment analysis.

Furthermore, we identified other classes such as complex SVs (n=148; 7.9%), reciprocal inversion (n=49; 2.6%; **Fig. 3a**), reciprocal translocation (n=30; 1.6%; **Fig. 3b**), and templated insertion (n=6; 0.7%; **Fig. 3c)**. The representative probands with simple SVs disrupting phenotype-relevant genes are shown in Figs. 3a-c. For example, we identified a templated insertion that disrupted *MECP2, w*hich has a well-established function in neurodevelopment^28^ in probands with NN disorders. This gene is known to be recurrently affected by dnSVs^10^ (**Fig. 3c**). This was independently validated by long-read sequencing (**Extended Data Fig. 2b**).

**Figure 3.**
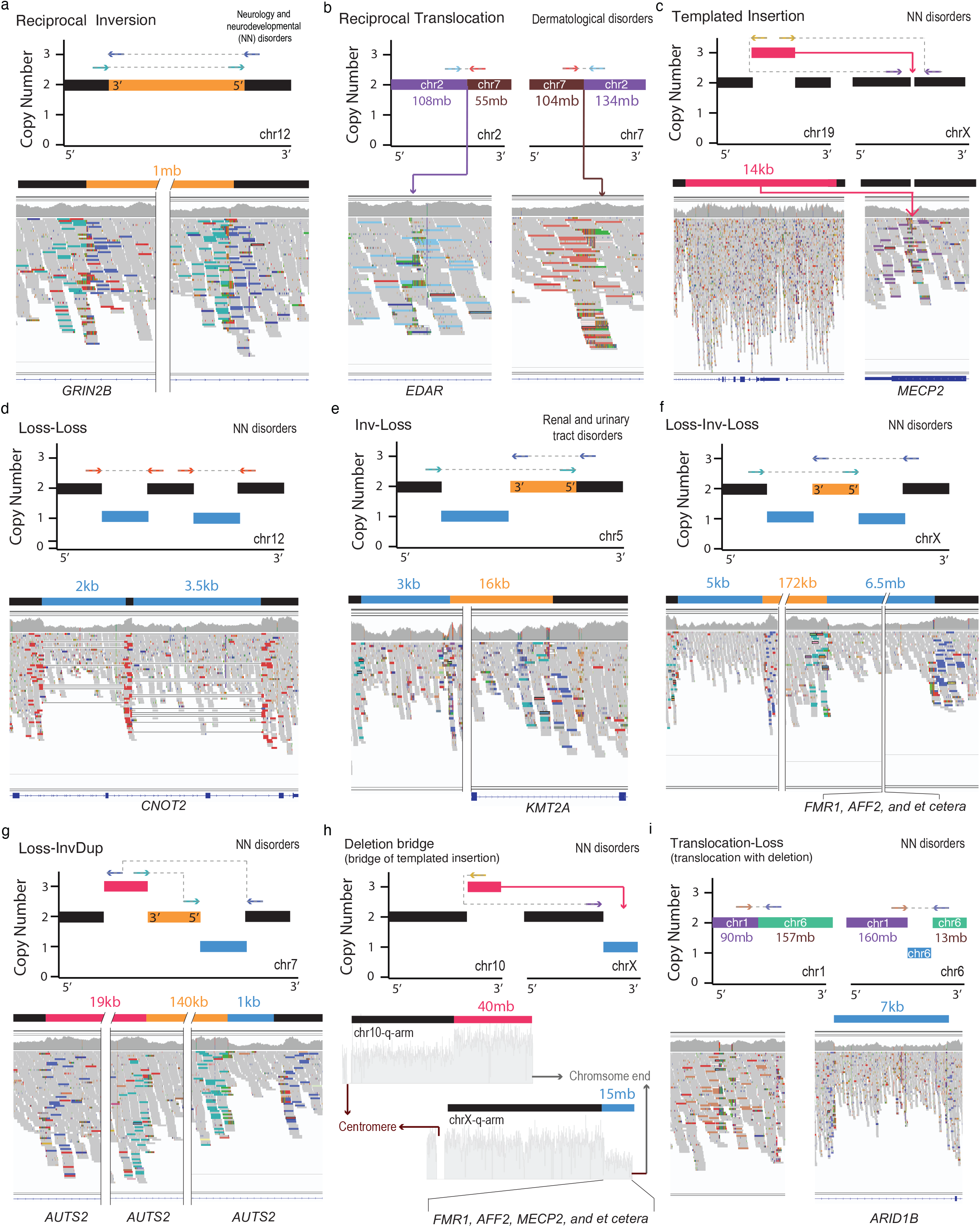
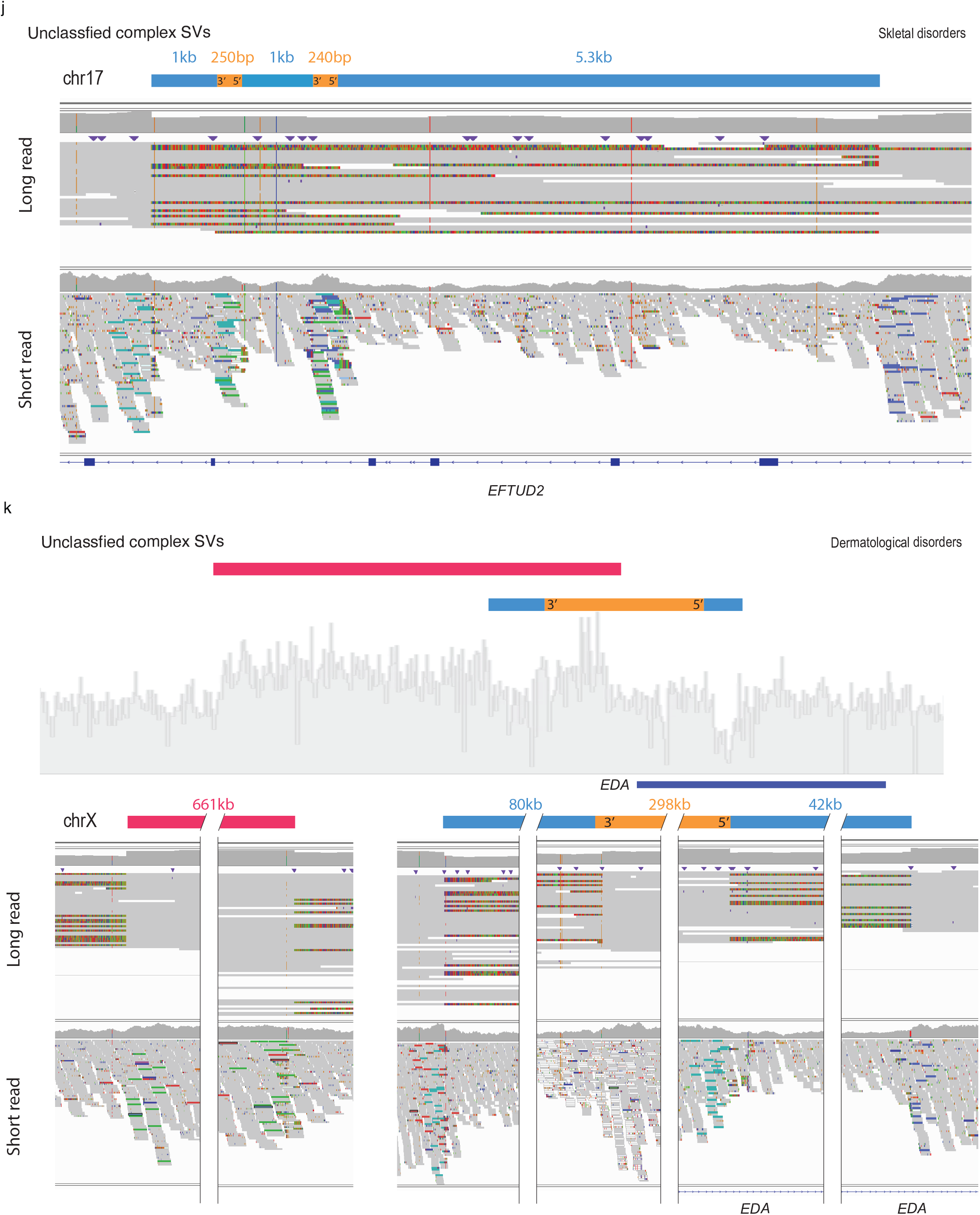
Representative simple and complex SVs disrupting potential causal genes. (**a-c**) simple SVs (**d-k**) complex SVs. (**a-i**) Schematic of major dnSV classes with copy numbers and read patterns (up). The schematic segments in blue, red, and orange represent deletion, duplication, and inversion, respectively. The size of the segments is not proportional to the SV size indicated above the segments. IGV screenshots illustrate dnSVs in probands (down). The potential causal genes affected by dnSVs are marked below the screenshots. (**j-k**) Unclassified types of complex dnSVs. The schematic segments (**j-k**) and coverage plot (k) are shown above IGV screenshots displaying long-read (up) and short-read (down) in probands.

We identified 41 cases of triplication dnSVs, which comprise 26 tandem duplications and 15 complex SVs (median = 574 kb; range 38 kb - 40 mb). We classified the timing of triplication of maternal origin into meiosis I and II (**Methods** and **Supplementary** Figure 1). This classification revealed that 83% of triplications in this cohort originated from maternal meiosis II (*P =* 1.51E-05; **Fig. 2d**). Further investigations using larger cohorts are required to confirm which step of meiosis contributes more significantly to dnSVs^29,30^.

### The role of complex SVs in rare disorders

Notably, the third most common type of dnSVs is complex SVs. Specifically, the proportion of complex SVs in our study (8%) is twice as high as in the previous most comprehensive dnSV study^24^ (4%; *P* = 4.00E-03; **Extended Data Fig. 3d**), highlighting that this SV type is an underestimated class of dnSVs in rare disorders. We further classified complex SVs into nine major classes (**Fig. 2c**). The most common class, termed ‘Loss-Loss’ (n=15; 10.1%), comprised two adjacent deletions (**Fig. 3d**). For instance, the two adjacent deletions (2 kb and 3 kb in length) in a proband with an NN disorder affected two exons in *CNOT2* (**Fig. 3d)** for which haploinsufficiency is known to cause a neurodevelopmental disorder with characteristic facial features^9^. In addition, adjacent deletions (5 kb and 1.7 kb in length) in a proband disrupted exon 2 of *AMER1* (**Extended Data Fig. 2c)** for which deficiency is associated with osteopathia striata with cranial sclerosis^31^. Other classes comprising inversion and deletion are ‘Inv-Loss’ (i.e., inversion with flanking deletion; n=14; 9.5%; **Fig. 3e**) and ‘Loss-Inv-Loss’ (i.e., paired deletion inversion; n=12; 8.1%; **Fig. 3f).** The representative cases with these types of complex SVs disrupting renal (and urinary) tract disorder-^32^ and NN disorder-related genes^33^ such as *KMT2A*, *AFF2*, and *FMR1*, are shown in Figs. 3e-f.

Another commonly observed class, termed ‘Loss-invDup’ (n=14; 9.5%), is characterised by copy-number loss plus nearby duplication linked by inverted rearrangements. For instance, a ‘Loss-invDup’ in a proband with NN affected an exon in *AUTS2*, implicated in neurodevelopment and as a candidate gene for numerous neurological disorders^34^. Another class, ‘Deletion bridge’ (i.e., bridge of templated insertion; n=7; **Fig. 3h** and **Extended data Figures 4a-b**), led to large deletions (720 kb in chromosome 1 in **Fig. 3h** and 15 Mb in chromosome X in **Fig. 3h**) containing genes involved in neurodevelopment^35,36^, such as *GALNT2*, *MECP2, and CTNNB1*, in probands with NN disorders (**Extended Data Fig. 4a**). ‘Translocation with deletion’ (n=3), led to deletions on either chromosome (**Fig. 3i**) or both chromosomes (**Extended Data Fig. 4c**), resulting in the disruption of phenotype-relevant genes such as *ARID1A*^37^ in a proband with NN disorders.

**Figure. 4.**
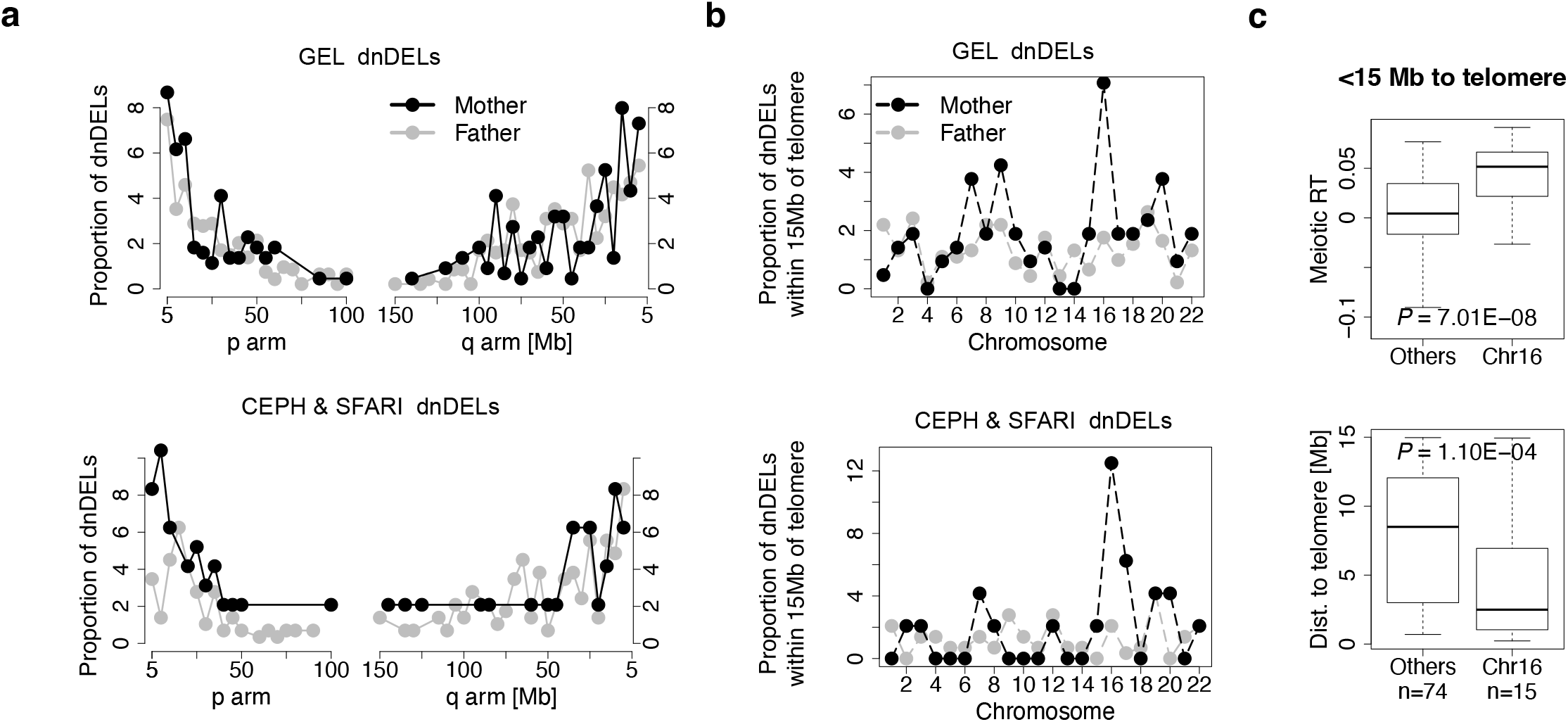
Enrichment of maternal dnDELs at subtelomeric, early-replicating regions of chromosome 16. (**a**) Proportion of dnDELs as a function of the distance to telomere ends (5 mb bins) in GEL (top) and in CEPH & SFARI cohort (bottom). (**b**) Proportion of dnDELs within 15 mb of telomere ends in GEL (top) and in CEPH & SFARI cohort (bottom). (**c**) Comparison of replication timing (top) and distance to telomere ends (bottom) of dnDELs between chromosome 16 and the other chromosomes.

The two remaining classes, comprising duplication and inversion, are ‘Dup-invDup’ (i.e., paired duplication inversion; n=7; 4.7%; **Supplementary** Figure 2) and ‘Dup-Trp-Dup’ (n=14; 9.5%; **Supplementary** Figure 2), exhibit structures involving two duplications linked by inverted rearrangements and duplication–inverted-triplication–duplication, respectively. Interestingly, beyond local-2 jumps (i.e., clusters of two rearrangements) as described above, we also found three instances of local-3 jumps involving three local rearrangements (**Extended Data** Figs. 5a). These are recurrently observed in human cancers^2^, while to our knowledge this is the first time that these events have been reported in the germline. The complex SVs that didn’t fit into the described classes were categorised as ‘Unclassified’ (n=59; 39.9%). Two of the complex SVs under the “Unclassified” category had long-read data that enabled us to resolve their genomic configuration (**Fig. 3j-k**). A proband with skeletal disorders (**Fig. 3j**) has a deletion-inversion-deletion-inversion-deletion structure which affected several exons in *EFTUD2* for which deficiency is likely to lead to craniofacial anomalies^38^. Another case has a structure of duplication followed by ‘Loss-Inv-Loss’ which disrupted the phenotype-relevant gene, *EDA*^39^ (**Fig. 3k**). In addition, a case with a small deletion (3 kb in chr22q.13.33) within a large deletion (20 kb in chr22q.13.33), where one of the breakpoints was the same, in a proband with an NN disorder disrupted *SHANK3* associated with a broad spectrum of neurodevelopmental disorders^40^ (validated by long-read in **Extended Data Fig. 5b**). Overall, our results highlight the complex nature of dnSVs in rare disorders.

### Clinical impact of dnSVs

Our analysis reveals that among probands with dnSVs, 9% (151/1,696) exhibit exon-disrupting pathogenic dnSVs associated with the probands’ phenotype. Notably, 12% (18/151) of these pathogenic dnSVs are complex SVs (**Fig. 2e**), emphasising the critical importance of thoroughly examining and considering complex dnSVs within the realm of rare disorders.

Furthermore, our investigation unveils distinctive patterns within intronic and intergenic dnSVs among probands with NN disorders. Intronic dnSVs showed a significant enrichment in known pathogenic genes associated with NN disorders in G2P database^41^ (*P* = 1.00E-03). In contrast, intergenic dnSVs, when assessed for genes within a 50 kb range up- and down-stream of the dnSVs (**Methods**), did not show such an association, suggesting the potential pathogenic role of intronic dnSVs in rare disorders (**Fig. 2f**). Additional studies using RNA-seq and/or CRISPR/Cas-9 genome editing are needed to elucidate the functional impact of these intronic dnSVs on mRNA splicing and expression.

### Genomic properties of dnSVs

In exploring genomic properties of dnSVs, we observed a prevalent distribution of *de novo* deletions (dnDELs) and tandem duplications (dnTDs) in gene-dense areas (**Extended Data** Fig. 6), in line with previous findings in somatic cells^2^. However, smaller dnDELs (<10 kb) are enriched in early-replicating regions (*P* = 1.00E-03), which is inconsistent with previous reports^2^.

We observed that the majority of dnSVs, primarily simple dnDELs, exhibit enrichment at the subtelomeric regions across autosomes (**Fig. 4a**). We also identified a positive association between the number of subtelomeric dnDELs and early replication regions, especially when they were within 15Mb of telomere ends (Spearman’s rho = 0.56, *P* = 6.46E-03; **Supplementary** Figure 3 and **Supplementary Table 1**). In total, the density of dnDELs within 15 Mb of telomere ends (i.e., 1.3/Mb) is 2.8 times greater than the autosome-wide average (i.e., 0.457/Mb).

Interestingly, we found that maternal dnDELs are enriched at the subtelomeric regions (<15 Mb) of chromosome 16 (**Fig. 4b**). These maternal dnDELs from chromosome 16 are not only replicated significantly earlier in meiosis than those from other autosomes (Wilcoxon rank-sum test, *P* = 7.01E-08; **Fig. 4c**), but also located significantly closer to the telomeres (*P* = 1.10E-04). In total, the density of maternal dnDELs within 15 Mb of telomere ends of chromosome 16 (i.e., 0.65/Mb) is 8.8 times greater than the maternal autosome-wide average (i.e., 0.074/Mb). We observed similar maternal enrichment of dnDELs in subtelomeric regions of chromosome 16 in an independent cohort^6^ (**Figs. 4a-b**).

## Discussion

Our investigation provides a substantial advancement in understanding of *de novo* structural variants in rare disorders, encompassing an extensive cohort of 13,702 parent–child trios. Particularly, we parent novel insights into the role of complex SVs with this domain thus far.

The prevalence of dnSVs, impacting 12% of probands, highlights the importance of integrating these variants into the broader spectrum of genetic factors contributing to rare disorders. Unlike conventional cytogenetic methods, such as array Comparative Genomic Hybridisation (CGH)-based technologies, WGS offers unparalleled precision in characterising the genomic configuration of complex dnSVs. This is particularly crucial as some simple deletions and insertions may be integral components of complex SVs often overlooked by array-based / exome-seq methods. For instance, in 37 cases where array and or exome-seq data were available, we found that 8 complex dnSVs (22%) were misclassified previously as simple dnSVs. In addition, 51 of 151 (34%) pathogenic SVs identified in our study were balanced rearrangements (e.g., balanced inversion) or CNVs affecting ≤ 2 exons that can’t be detected by array-based / exome-seq methods, highlighting the importance of WGS-based routine genetic screening.

Notably, dnSVs exhibited non-random distribution patterns, showing enrichment in specific genomic locations associated with distinct features depending on the parent of origin. Strikingly, we observed an enrichment of maternal dnDELs within 15 Mb of the telomeres of chromosome 16. This enrichment positively correlates with skewed early replication regions across chromosomes. While the genomic basis for this maternal bias in subtelomeric regions of chromosome 16 is unknown, previous reports have suggested potential explanations^42^. These include early subtelomeric replication in meiosis^43^, increased rates of meiotic double-strand breaks in the distal parts of chromosome^44^, and or biased maternal non-crossover gene conversion^45^. Overall, these findings indicate the need for further investigation into parental influence and region-specific impacts on disease manifestation.

We note several limitations, which open potential venues for future investigations. Due to the lower detection sensitivity of *de novo* retrotransposition, with the current pipeline, these variants have been excluded from the analysis. Specifically, we observed a rate of 0.01 events per genome which is far lower than expected (0.03-0.038 per genome^24,46^). Furthermore, the limitations of this dataset constrained CNV detection to read-based algorithms, which, compared to read alignment patterns algorithm, have lower accuracy. We estimated our false negative rate to be only 6% using read-pattern-based callable SVs (i.e., Manta), but this increased to 20% when including CNVs called only by read-depth algorithms (identified by array in the DDD project^21^ and validated by CANVAS in our WGS data). Future research could explore techniques such as long-read sequencing^47^ to enhance our ability to detect dnSVs in repetitive regions^48^ and CNVs. Furthermore, the identification of inherited pathogenic SVs will be needed to facilitate a more comprehensive diagnosis.

Overall, our findings expand the understanding of dnSVs in rare disorders and highlight the need for ongoing research to unravel the nuance of their contribution, diagnostic challenges, and potential clinical applications.

## Online Methods

### SV calling

The cohort was whole-genome sequenced and read alignment and SV calling using Isaac^49^ and Manta^20^, respectively, were performed by the Genomics England Bioinformatics team^19^. The details of sequencing and variant calling have been previously described. Manta VCF files were converted to BEDPE format using SVtools^50^ and then BEDTools^51^ was utilised to extract proband-specific SVs with ≥ 50 bp in length for each family (**Supplementary** Figure 4). We removed SVs having evidence of clipped reads (i.e., split reads) at breakpoints in either parent. Specifically, SVs supported by ≥ 4 clipped reads at either breakpoint or ≥ 2 clipped reads at both breakpoints in either parent were excluded. The genomic coordinates of SVs called from hg19-aligned genomes (1915 probands) were converted to hg38 genomic coordinates using LiftOver^52^. SVs found in 3+ samples were removed because such SVs were likely germline SVs and artifacts. We selected SVs flagged as “PASS” or “MGE10kb” and further narrowed down SVs with the Manta score > 30 and supporting discordant reads > 10. We rescued SVs with imprecise breakpoints if they were supported by CANVAS^53^. We excluded SVs with VAF < 0.1 to remove mosaic SVs. Translocation through retrotransposon-mediated 3′ transduction was excluded to focus on dnSVs. All SVs were manually validated to identify high-confident de novo SVs using IGV browser^54^. Long-read sequencing data and diagnostic SNV/Indels (**Fig. 1a)** were obtained from Genomic England^19^. Insertion events called by Manta^20^ were further classified into retrotranspositions using RepeatMasker. This process resulted in a lower sensitivity because Manta with Issac-based alignment is not optimized to call retrotranspositions. Retrotransposition-specific identification tools, such as MELT^55^ or xTea^56^, are needed to increase sensitivity for retrotransposition detection.

### SV classification

We used ClusterSV^2^ (https://github.com/cancerit/ClusterSV) to group rearrangements (i.e., breakpoints) into rearrangement clusters. We defined complex SVs as those with ≥ 2 clustered breakpoints except for simple SVs involving reciprocal inversion, balanced translocation, templated insertion, and dispersed duplication. In general, we classified the types of complex SVs according to the previous study that comprehensively characterised somatic complex SVs using thousands of cancer genomes^2^. In short, complex SVs involving two inversions were categorised into Loss-invDup, Dup-Trp-Dup, Inv-Loss (i.e., inversion with flanking deletion), Loss-invLoss (i.e., paired deletion inversion), and Dup-invDup (i.e., paired duplication inversion) according to read patterns and copy numbers (**Supplementary** Figure 2). Complex SVs involving two deletions were classified as Multi-Loss. Bridge deletion (i.e., bridge of templated insertion) and Translocation-Loss (i.e., translocation with deletion) were classified using the previously described criteria^2,57^. Local-3 jumps involving three local rearrangements were discovered according to the read patterns described in the previous cancer study^2^. Breakpoints filtered out near unresolved SV classes were rescued if they could resolve the unresolved SV classes according to the types of SV defined. Complex SVs that didn’t fit into the described classes were classified as ‘Unclassfied’. All complex SVs were manually validated using IGV browser^54^, Samplot^58^, or BamSnap^59^.

### SV phasing to identify parent of origin and estimate the timing of triplication from maternal origin

We used unfazed^60^, which employs both read-based phasing and SNV allele-balance phasing, to identify parent of origin for dnSVs. To classify the timing of maternally-derived triplication into meiosis I and II, we first identified triplications (including those in complex SVs) from maternal origin (step 1) and further classified them into meiosis I and II (step 2) using a set of informative genotypes (**Supplementary** Figure 1). At least five SNPs were required for each step.

### Evaluation of clinical relevance of dnSVs

The identified SVs disrupting exons were reviewed for potential clinical relevance by NHS clinical scientists and/or Genomics England. We considered SVs as being potential (likely) pathogenic SVs if at least one of the following criteria were fulfilled: i) the variant had been clinically assessed as likely pathogenic or pathogenic by an NHS genomic laboratory hub ii) the variant had been reviewed on a research basis and considered appropriate to return through the Diagnostic Discovery pathway to an NHS genomic laboratory hub for evaluation.

### Enrichment test of non-coding dnSVs in known pathogenic genes in NN disorders

We first extracted the non-coding dnSVs (i.e., intronic and intergenic dnSVs) for which genomic coordinates didn’t include any exons in NN disorders based on the Gencode basic V45 GTF file and then obtained the known pathogenic genes associated with NN disorders from the Gene2Phenotype developmental disorders panel^41^. Specifically, we kept all genes with organ specificity equal to “Brain/Cognition”, allelic requirement equal to “monoallelic_autosomal”, and a confidence category equal to “strong” or “definitive” (n=190 genes). We then computed the observed-over-expected ratio for the overlap between the non-coding dnSVs and known pathogenic genes in NN disorders using the Genome Association Tester software^61^. Intronic and intergenic regions were obtained based on the canonical transcript of protein-coding genes in the Gencode basic V45 GTF file using BioMart and GencoDymo R packages, and bedtools^51^. These two regions were used as a workspace in GAT to test the over-representation of intronic and intergenic dnSVs in intronic and intergenic regions of the known pathogenic genes, respectively. We added a window of 5-500kb (5, 10, 25, 50, and 500kb) up- and down-stream to each intergenic dnSVs to perform the enrichment test. Number of random samples (“--num-samples”) for each GAT run was set to 1000.

### Distribution of dnSVs across genomic properties

Genomic property metrics and fragile site (FS) information were downloaded from PCAWG structural variation paper based on hg18^2^. Only 1,362 out of 1,377 GEL de novo simple deletions were able to liftover from hg38 to hg18 for further association analysis with the genomic properties. Telomere (chromInfo.txt.gz) and centromere (cytoBand.txt.gz) information were downloaded from UCSC Genome Browser (http://hgdownload.cse.ucsc.edu/goldenPath/hg19/database/). To test for associations between dnSVs and the above library of genomic properties, the median of real dnSV positions (one random side from each breakpoint were chosen to reduce dependence) were compared to 1,000 median random positions using Monte Carlo simulation method. For each genomic property, the frequencies of random medians generated by the simulation will form a normal distribution. The empirical p values were calculated by the sum of random medians lower or higher than real median, then divided by 1,000. To calculate the median shift, the real observations from each genomic property were normalised on a scale from 0 to 1. Then the the shift of the median are calculated by the difference between the observed median and 0.5, assuming a uniform distribution has a median of 0.5

### Enrichments near telomeres and centromeres

We equally partitioned the genome into 5-Mb bins based on their distance to the telomere ends. For comparison, we also partitioned the genome based on their distance to the centromeres. We identified that dnSVs are enriched near the telomere ends. However, the dnSVs are more evenly distributed to the centromeres. For the validation cohort, we downloaded the all_dnsv.csv file from Belyeu et al. 2021^24^. In total, there are n=309 CEPH and SFAI dnSVs across autosomes after removing chrX and chrY. And finally, only n=192 dnDELs were used in validation analysis.

### Replication timing profiles

The meiotic replication timing (RT) profile was downloaded from Pratto et al., 2021^43^ using table expRT_MeiS_ALL_S2_2to4C_SCP3_DMRT1, which was profiled from S-phase nuclei sorted from human testis. The reference mitotic RT was downloaded from Koren et al., 2014^62^, which was profiled from unsynchronised, flow-sorted lymphoblastoid cell lines. The Replication timing skew (RTS) values were calculated as the difference between the proportions of early (E) and late (L) replication regions per chromosome as:

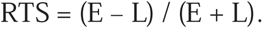

## Code availability

Custom Python and R scripts for data analysis can be found at https://github.com/hj6-sanger/GEL_SV

## Acknowledgments

We thank the families and their clinicians for their participation and engagement, and our colleagues who assisted in the generation and processing of data. We would like to thank Ana Lisa Taylor Tavarer for helpful discussions and advice. This research was made possible through access to the data and findings generated by the 100,000 Genomes Project. The 100,000 Genomes Project is managed by Genomics England Limited (a wholly owned company of the Department of Health and Social Care). The 100,000 Genomes Project is funded by the National Institute for Health Research and NHS England. The Wellcome Trust, Cancer Research UK and the Medical Research Council have also funded research infrastructure. The 100,000 Genomes Project uses data provided by patients and collected by the National Health Service as part of their care and support. This research was supported in part by the Wellcome Trust grant.

## Author Contributions

R.R. conceived the project. R.R., P.C., and M.H. supervised the project. H.J., T.Y., and R.R. wrote the manuscript; all authors reviewed and edited the manuscript. H.J., T.Y., led the analysis of the data with help from P.D., I.G.S., M.D.C.N. S.W. reviewed the pathogenic complex dnSV candidates. H.F., helped with clinical interpretation of the dnSVs. A.S., M.H., P.C., H.J., and R.R. helped with data interpretation and statistical analysis.

## Extended data Figure legend

**Extended Data Fig. 1.** dnSV rate (**a**) Proportion of samples by rare disorder types (**b**) Comparison of the rate of dnSVs between our study and previous three studies. The confidence interval was computed using the Exact binomial test. (**c**) dnSV rate across rare disorder types. Rare disorder types having ≥ 300 samples except for unaffected siblings are shown.

**Extended Data Fig. 2.** Validation of simple (a-b) and complex dnSVs (c) by long-read.

**Extended Data Fig. 3.** Analysis of dnSV (**a**) Correlation analysis between dnS and dnS SNV/indel burden using all (up) and NN samples (down). The P values were determined based on Spearman correlation analysis (**b**) Comparison of maternal and paternal age at birth between probands with dnSVs and those without dnSVs in Dysmorphic and Congenital abnormality syndromes and skeletal disorders. The P values were calculated based on a one-sided t-test and were corrected using the Benjamini-Hochberg method. (**c**) Distribution of the size of simple deletion and tandem duplication. (**d**) Comparison of the size of dnDELs according to parent of origin in CEPH & SFARI cohort. (**e**) Comparison of the proportion of complex de novo SVs between our and the previous study. The mobile element insertions in the previous study were not included. (**f**) Distribution of the length of microhomology at breakpoint junctions.

**Extended Data Fig. 4.** Representative cases with ‘Translocation-Loss’. Two subclasses of ‘Translocation-Loss’: ‘Deletion bridge’ (**a**) and ‘Translocation with deletion’ (**b-c**). (**a**) An additional small deletion (size=2kb) in chr19 was found in the templated sequence. (**b-c**) Translocations accompanying deletions on either chromosome (**b**) or on both chromosomes are shown.

**Extended Data Fig. 5.** Representative cases with complex SVs. (**a**) Local-3-jumps (**b**) Unclassified complex SV type

**Extended Data Fig. 6.** Genomic properties of dnSVs. **(a-b)** Distribution of dnDELs (a) and de novo tandem duplications (b) in the context of gene density (top) and replication timing (bottom). P values were calculated using Monte Carlo simulation test (\**P* < 1E-03). Each density plot is coloured by the extent of the median shift below (blue) or above (red) 0.5, assuming that a uniform distribution has a median of 0.5.

