## Extended_Data_Figures for "Deciphering the role of germline complex *de novo* structural variations in rare disorders"

Extended Data Figure 1

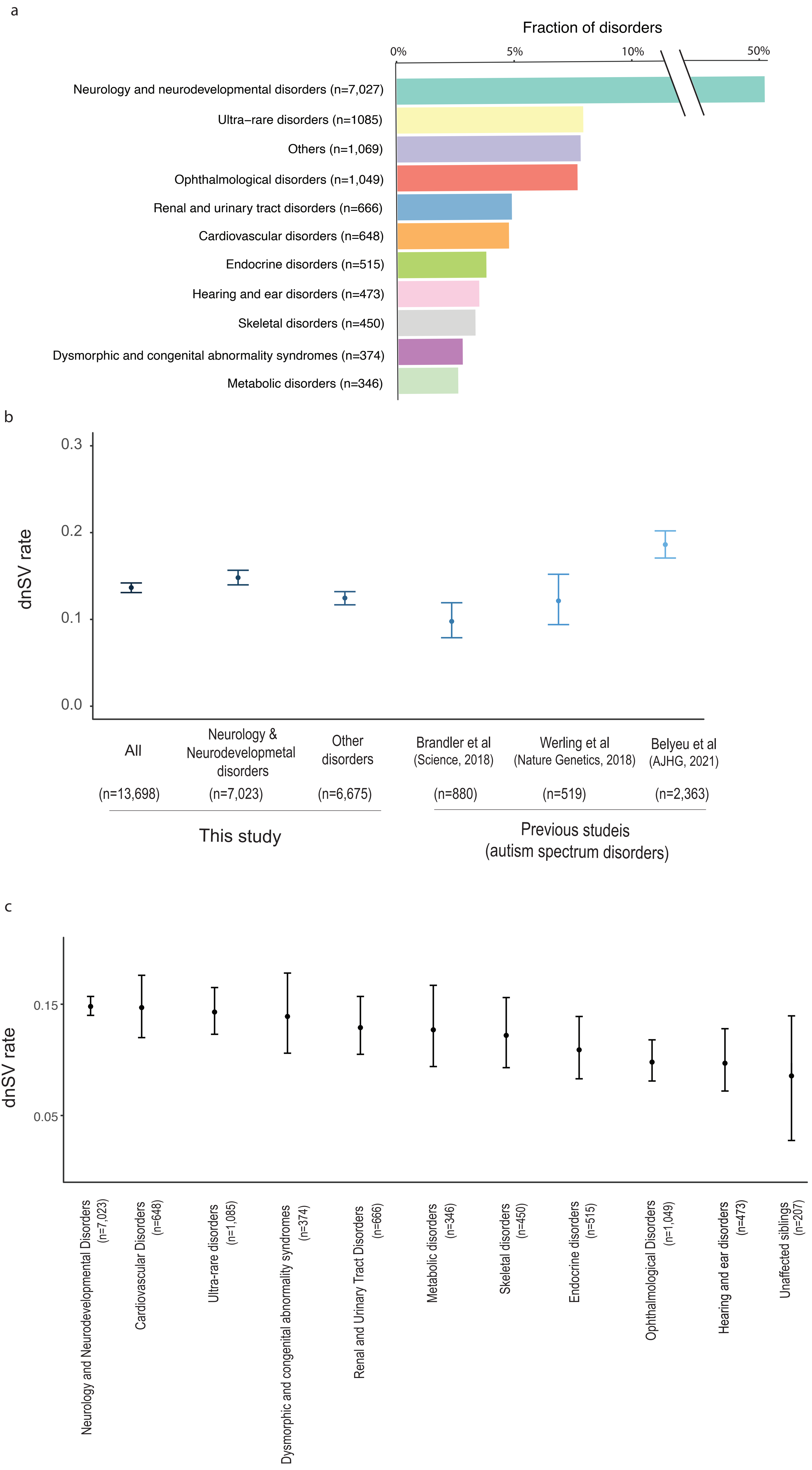

Extended Data Figure 2

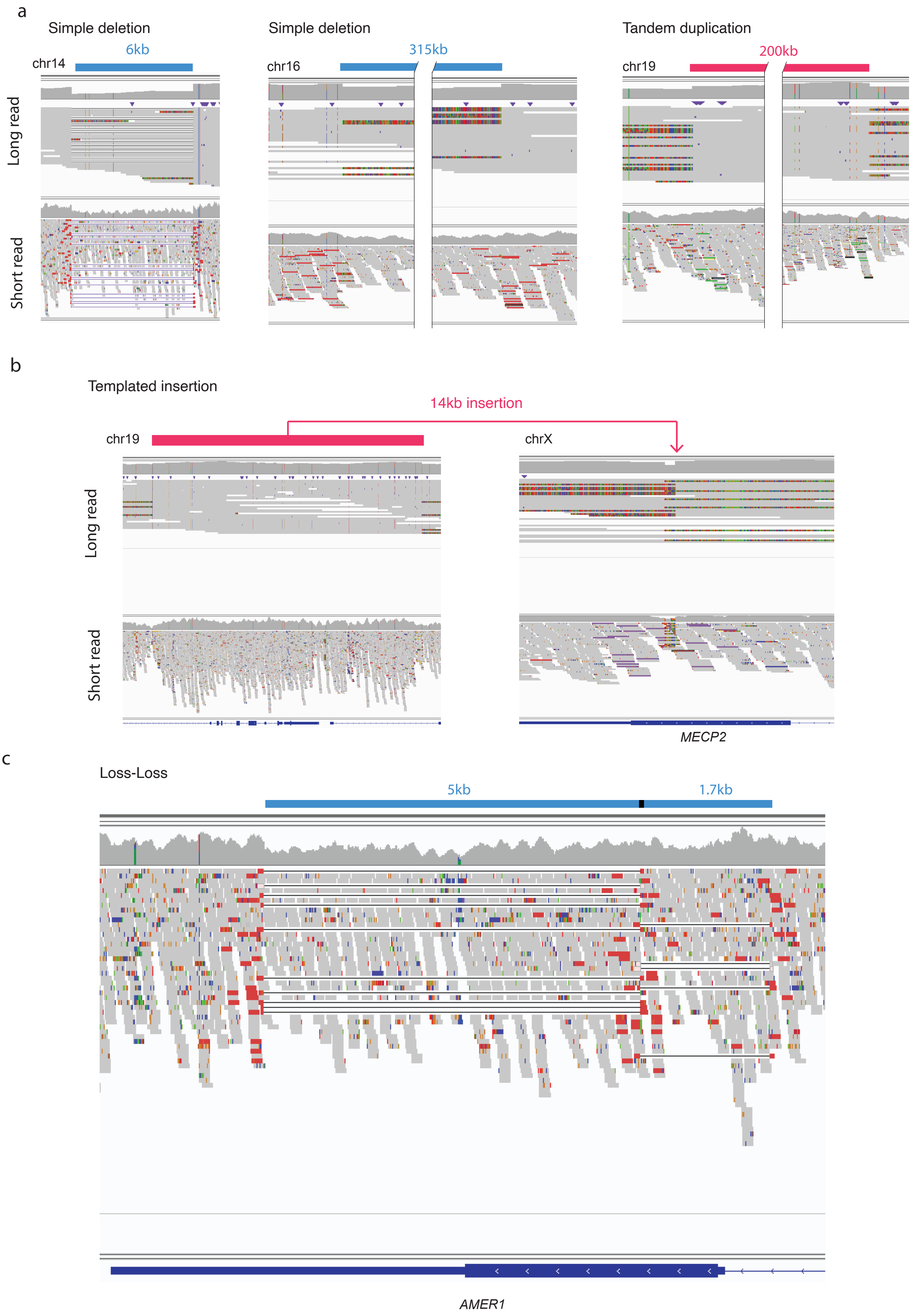

Extended Data Figure 3

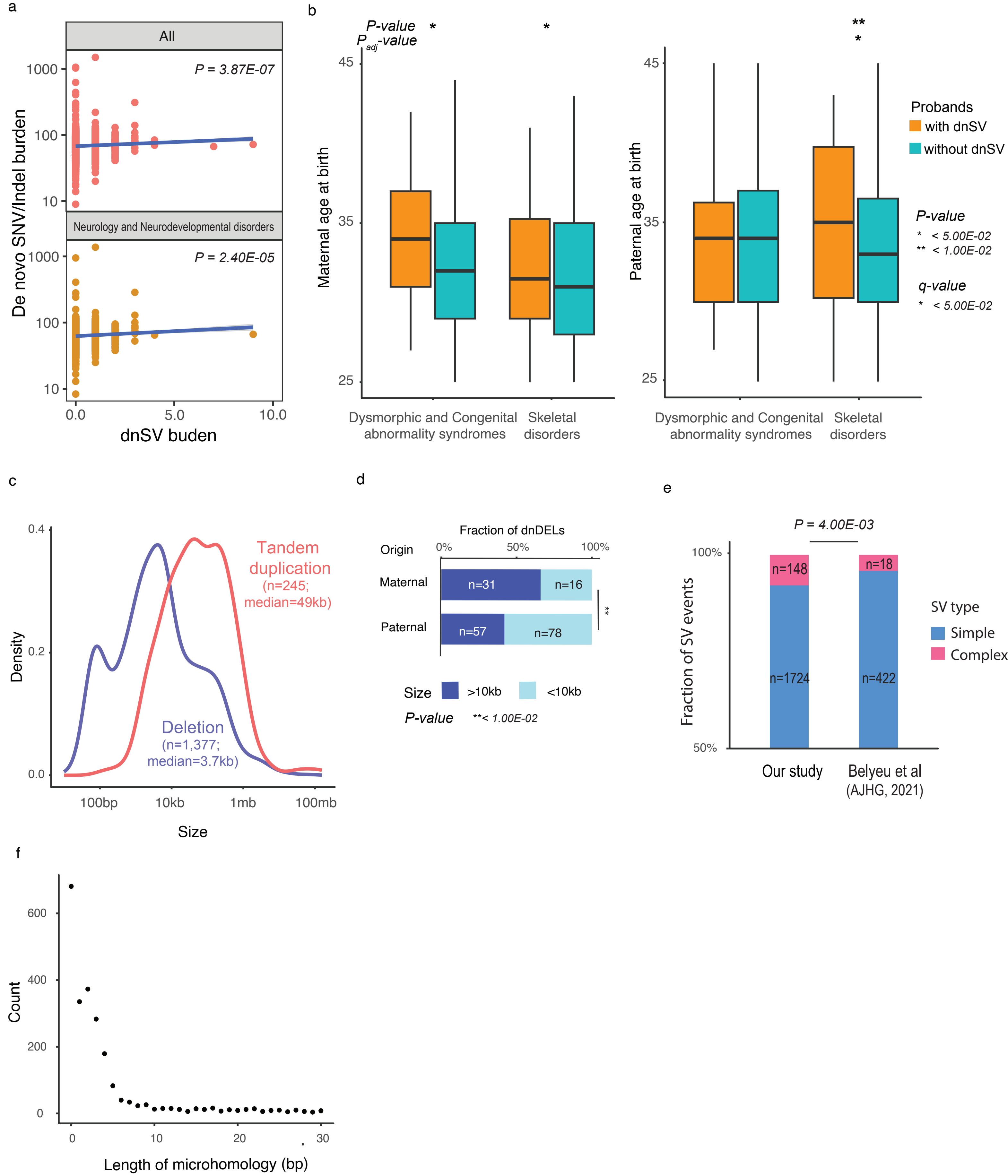

### Extended Data Figure 4

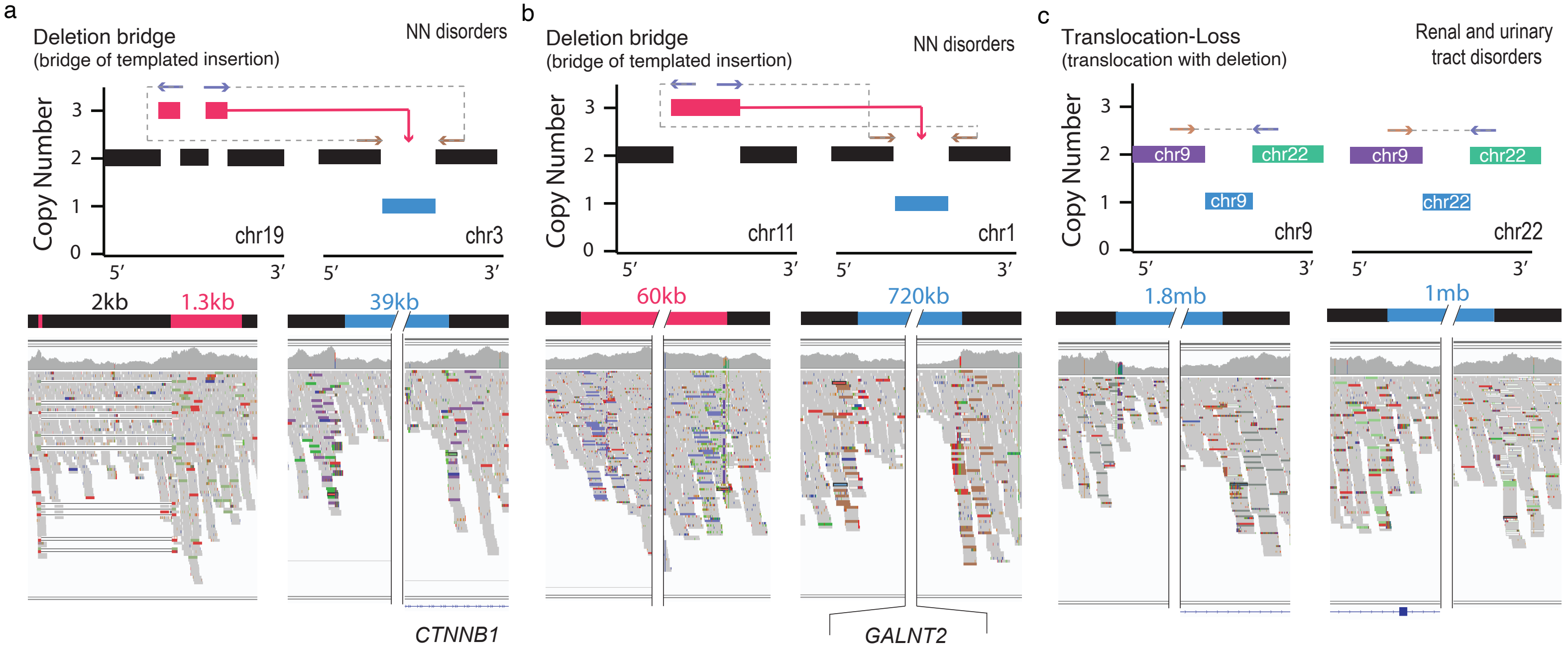

Extended Data Figure 5

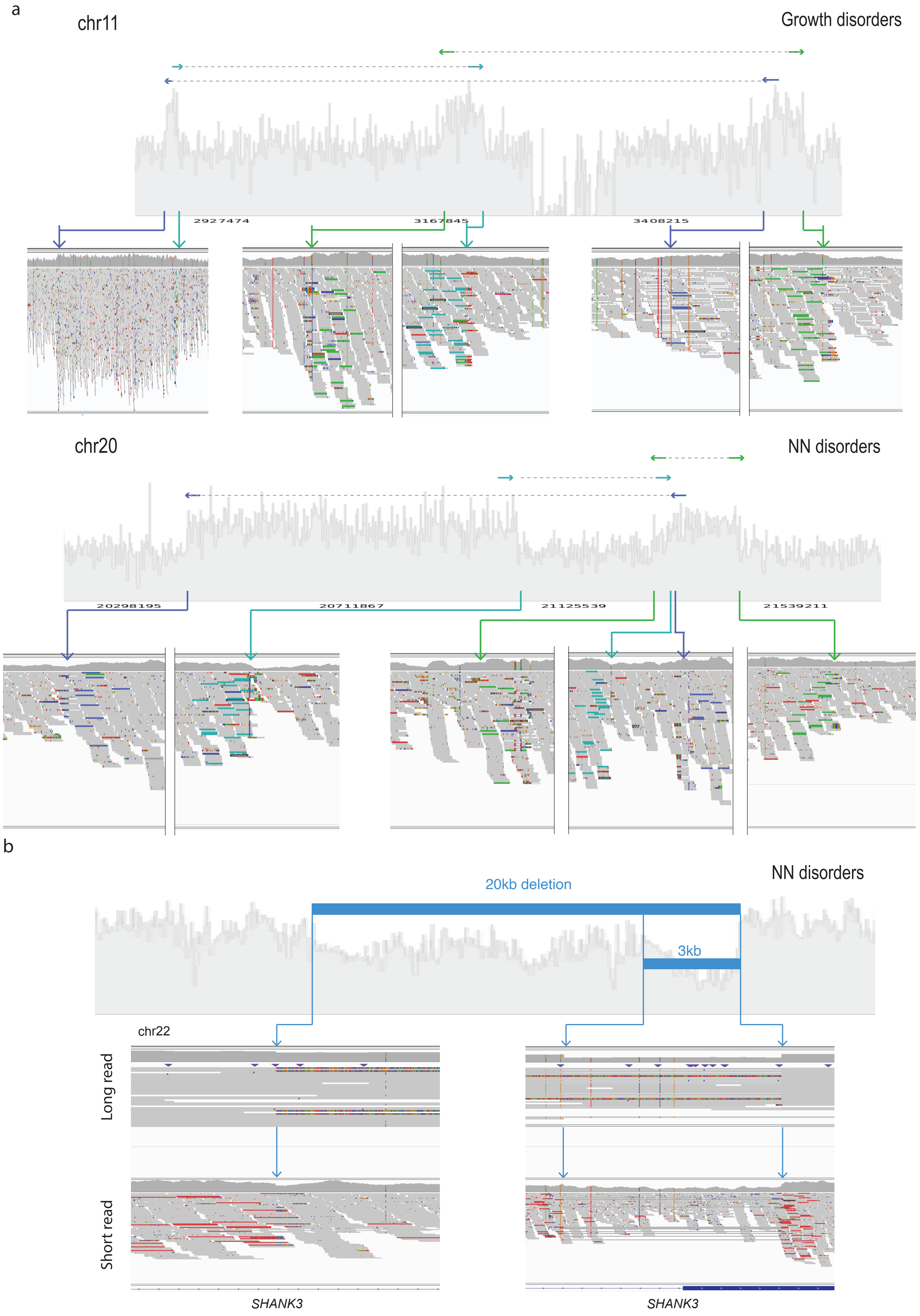

Extended Data Figure 6

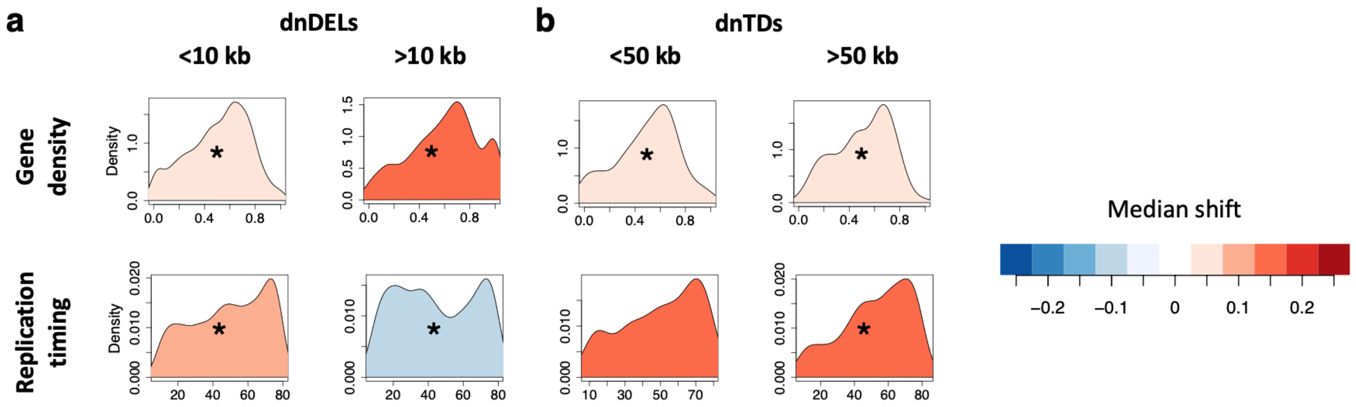
